# A mechanism coordinating root elongation, endodermal differentiation, redox homeostasis and response

**DOI:** 10.1101/813774

**Authors:** Jing Fu, Jiaming Liu, Xudong Gao, Xinglin Zhang, Juan Bai, Yueling Hao, Hongchang Cui

## Abstract

Root growth relies on both cell division and elongation, which occur in the meristem and elongation zones respectively. SCARECROW (SCR) is a GRAS family gene essential for root growth and radial patterning in the Arabidopsis root. Previous studies showed that SCR promotes root growth by suppressing cytokinin response in the meristem, but there is also evidence that SCR expressed beyond the meristem is required as well for root growth. Here we report that SCR promotes root growth by promoting cell elongation through suppression of oxidative stress response and maintenance of redox homeostasis in the elongation zone. In the *scr* root, a higher level of hydrogen peroxide was detected, which can be attributed to down-regulation of peroxidase gene 3. When stress response was blocked or redox status was ameliorated by the *aba2* or *upb1* mutation, the *scr* mutant produced a significantly longer root with longer cells and a larger and mitotically more active meristem, even though the stem cell and radial patterning defects still persisted. We showed that *WRKY15*, an oxidative responsive gene, was a direct target of SCR down-regulated in the *scr* mutant, which suggests that SCR has an active role in suppressing oxidative stress response. Since hydrogen peroxide and peroxidases are essential for endodermal differentiation, these results suggest that SCR plays a central role in coordinating cell elongation, endodermal differentiation, redox homeostasis, and oxidative stress response in plant root.

**One sentence summary:** This study reveals a novel mechanism of root growth regulation, which involves a previously unrecognized role of SCR in regulating cell elongation, endodermal differentiation, and redox homeostasis.

## Introduction

As the major organ for nutrient and water uptake, the root system must expand its surface area continuously to meet the need of a growing plant. Continuous growth of the root relies on the mitotic activity of the root apical meristem (RAM) and the maintenance of a functional quiescent center (QC), which is the source of all cells in the root and is hence regarded as the stem cells of the root. However, root length is determined by not only the number of cells but also the final cell size.

An essential role for redox homeostasis in stem cell renewal in plants has been established (Tang et al., 2017). While the QC and the RAM are associated with an oxidized environment (Jiang et al., 2003; Takahashi et al., 2004), the elongation zone is more reduced (Tsukagoshi, 2015). When the redox potential is altered in the RAM, such as in the Arabidopsis *rml1* or *app1* mutant, the RAM is lost or severely reduced (Del Pozo, 2016; Vernoux et al., 2000). The redox status in the elongation zone also affects root meristem activity, as the RAM becomes larger in the *upb1* mutant due to a more reduced redox status in the elongation zone (Tsukagoshi et al., 2010). When the redox status in the elongation zone is more oxidized, cell elongation is inhibited, resulting in a short root phenotype (Mabuchi et al., 2018; Tsukagoshi, 2015).

Reactive oxygen species (ROS), including hydrogen peroxide, oxygen superoxide, and free oxygen radicals, are normally produced at a low level as a byproduct of the energy generating process in mitochondria, the photosynthesis reaction in chloroplasts, and the photorespiration in peroxisomes, but their levels can be elevated when growth conditions are suboptimal (Choudhury et al., 2017). ROS can also be generated by active mechanisms in some cell types, such as lateral root, root hair, and xylem, where they play a positive role in the cell differentiation programs (Bagniewska-Zadworna et al., 2014; Gapper and Dolan, 2006; Mangano et al., 2016; Orman-Ligeza et al., 2016). In the endodermis, for example, hydrogen peroxide is required for the formation of the Casparian strip, a structure essential to this cell type’s function in selective nutrient uptake and defense against pathogens (Dinneny, 2014; Robbins et al., 2014). Accordingly, many genes involved in hydrogen peroxide formation and catabolism, such as Reactive Oxygen Burst Homolog (ROBH) and peroxidase genes, are highly expressed in the endodermis (Lee et al., 2013). Conceivably, if these genes are expressed abnormally, hydrogen peroxide could accumulate, which in turn would affect root growth. How these antagonizing processes are coordinated to ensure normal growth and development is currently unknown.

SCARECROW and SHORT-ROOT (SHR) are key regulators of root growth and development in *Arabidopsis thaliana*. In the *scr* and *shr* mutants, the QC is lost, consequently resulting in the loss of the RAM and cessation of root growth (Di Laurenzio et al., 1996; Helariutta et al., 2000). The *scr* and *shr* mutants also have a radial patterning defect, which is characterized by the loss of a cell layer due to the absence of an asymmetric cell division that gives rises to the endodermis and cortex (Di Laurenzio et al., 1996; Helariutta et al., 2000). As expected, SCR is expressed in the QC, the cortex/endodermis initial cell, and the endodermis (Di Laurenzio et al., 1996). In contrast, SHR is expressed in the central stele (Helariutta et al., 2000). However, the SHR protein is able to move into the neighboring cells, including the QC and endodermis (Nakajima et al., 2001), where it forms a heterodimer with SCR (Cui et al., 2007). The SHR-SCR complex in turn activates genes that are involved in cell-cycle and cell-fate determination (Cui et al., 2007; Moreno-Risueno et al., 2015; Sozzani et al., 2010). The SHR-SCR complex also enhances SCR expression through a positive feedback loop, leading to SCR accumulation (Cui et al., 2007). The superfluous SCR protein not only blocks SHR movements but also has an SHR-independent role in sugar response and radial patterning (Cui and Benfey, 2009a, b; Cui et al., 2012; Cui et al., 2007).

Previous studies have demonstrated a crucial role for QC-expressed SHR and SCR in stem cell renewal and root growth (Sabatini et al., 2003). However, there is also evidence that SHR and SCR expressed in the elongation zone also contribute significantly to root growth (Moubayidin et al., 2016; Sabatini et al., 2003; Sebastian et al., 2015). To elucidate the mechanism by which SCR regulates root growth and development, we have identified SHR and SCR direct targets at the genome scale by ChIP-chip (Cui et al., 2011; Iyer-Pascuzzi et al., 2011; Sozzani et al., 2010). Intriguingly, we found that many SHR and SCR target genes are associated with abiotic stress. In further studies, we showed that the *scr* mutant is hypersensitive to ABA, suggesting that SCR may play a role in abiotic stress response (Cui et al., 2012). In the present study, we explored the role of SCR in abiotic stress and its relationship to root growth, which led to the finding that SCR promotes root cell elongation by mitigating oxidative stress response as well as maintaining a more reduced cellular redox status.

## Results

### The root growth defects in *scr* and *shr* are alleviated by blockage of ABA signaling

Previously we have shown that the *scr* mutant is hypersensitive to ABA (Cui et al., 2012). Since ABA is a hormone associated with abiotic stress, this finding raises the interesting possibility that the root growth defect of the *scr* mutant is due, at least partially, to an abnormal level of abiotic stress or stress response. If this is true, blocking ABA biosynthesis should alleviate the short root phenotype in *scr*. To test this hypothesis, we generated a double mutant with *scr-1* and the ABA deficient mutant *aba2-1/gin1-3*. As expected, we indeed found that the *scr-1 aba2-1* double mutant had significantly longer roots than the *scr-1* single mutant (Fig 1A and B). Hence, we have uncovered a new role of SCR in promoting root growth through suppression of abiotic stress response.

**Fig 1.**
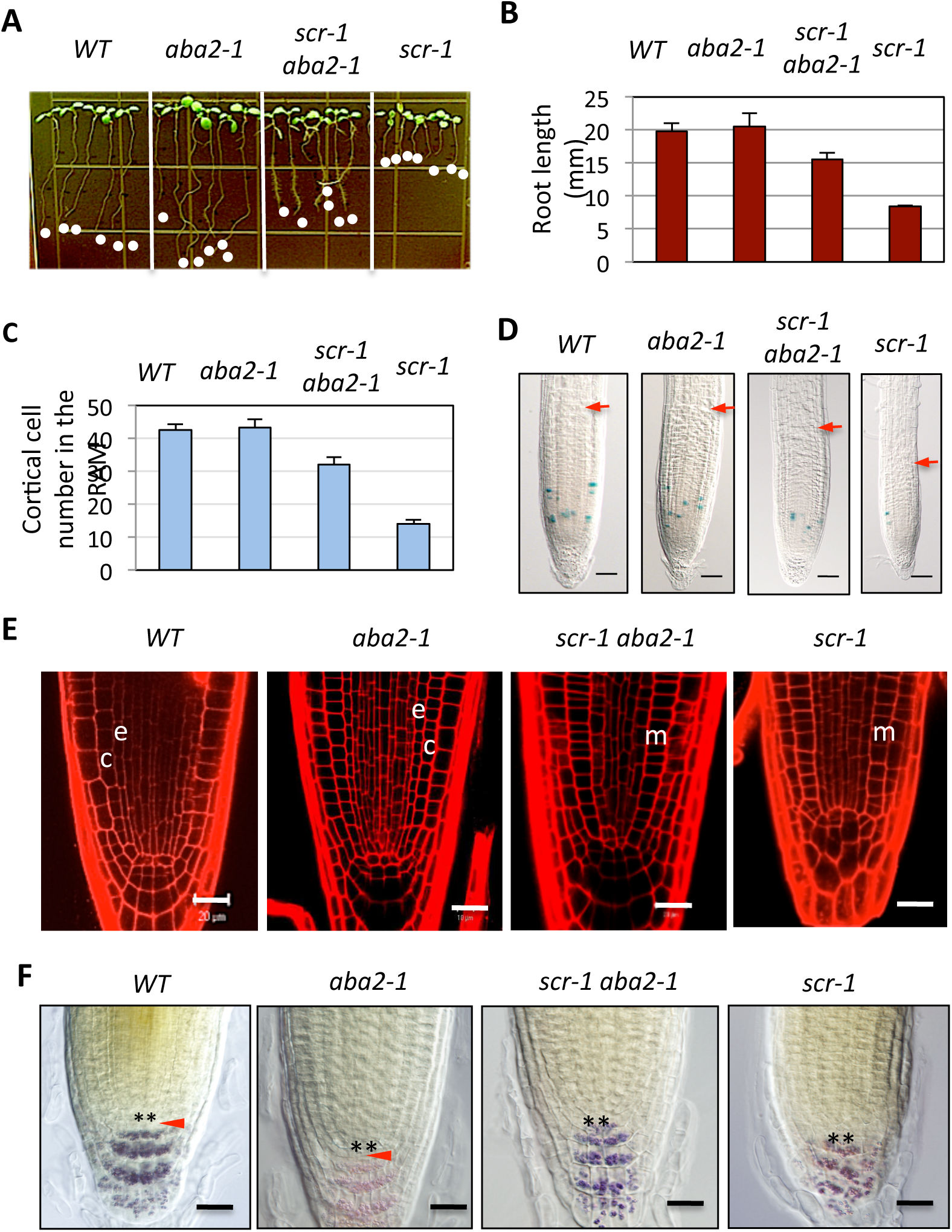
Blocking ABA signaling alleviates root growth phenotype in *scr* mutant, but not QC and radial patterning defects. A. Representative image showing seedlings grown in MS medium 8 days after germination. *WT*, wild type. B-C, Root length and cortical cell number in the RAM, respectively. Data are represented as mean ± SEM. N>20. D. Light microscopy images showing meristem activity. Blue spots are derived from the *pCYCB1;1-GUS* reporter construct, indicating cells the *G2*-M phase *o*f the cell cycle. Bars, 50 µm. E. Confocal microscopy images showing the radial patterning. e, endodermis; c, cortex; m, mutant cell layer. Bars, 20 µm F. Starching staining. Marked by the red arrows and asterisks are columella stem cells and QC respectively. Bars, 20 µm.

Since SHR and SCR act in the same pathway in root growth and development (Petricka et al., 2012), we also generated the *shr-2 aba2-1* double mutant and examined the effect of the *aba2* mutation on root growth of the *shr* mutant. Although the *shr-2 aba2-1* root was also longer than that of the *shr-2* mutant (Fig S1A and B), the improvement was much less than for the *scr-1 aba2-1* double mutant. This is not totally surprising, as previously we have shown that SHR and SCR have distinct roles in radial patterning (Cui and Benfey, 2009b; Cui et al., 2011), cytokinin homeostasis (Cui et al., 2011), and sugar signaling (Cui et al., 2012).

### The *aba2* mutant enhanced RAM activity, but does not rescue the radial patterning and stem cell defect, in the *scr* and *shr* mutants

The longer root of the *scr-1 aba2-1* double mutant could result from a larger root meristem, higher mitotic activity, rescued stem cells and/or radial patterning. To distinguish among these possibilities, we first examined the RAM in the *scr-1 aba2-1* double mutant, the *scr-1* single mutants, and the wild type. As shown in Fig 1C, the *scr-1 aba2-1* double mutant clearly had a larger meristem that the *scr-1* single mutant. In line with this observation, the *scr-1 aba2-1* double mutant also had a greater number of cortical cells in the meristem zone (Fig 1C). To examine mitotic activity in the RAM, we introduced the *CYCB1;1:GUS* reporter gene into the different mutant backgrounds. A relatively larger number of cells with GUS activity (Fig 1D) were visible in the *scr-1 aba2-1* double mutant, indicating a mitotically more active meristem.

To determine whether the radial patterning and stem cell defects in the *scr* mutant were rescued in the *scr-1 aba2-1* double mutant, we examined the cell organization by confocal microscopy. Interestingly, the *scr-1 aba2-1* double mutant still had only a single layer of ground tissue characteristic of the *scr* mutant (Fig 1E). The cells at the QC position in the *scr-1 aba2-1* double mutant were still disorganized, suggesting that the stem cell defect was not rescued. To further determine the properties of the cells at the QC position, next we examined the presence of starch granules by iodine staining. In wild type and the *aba-2* mutant, a starch-free cell layer known as the columella stem cells lies beneath the QC (Fig 1F). In the *scr-1* single mutant and the *scr-1 aba2-1* double mutant, the cells at the corresponding position contain starch grains, indicating the loss of stem-cell property (Fig 1F). Similar results were obtained with the *shr-2 aba2-1* double mutants (Fig S1, C-E). These results suggest that the improved root growth of the *scr-1 aba2-1* and *shr-2 aba2-1* mutants can be attributed to enhanced RAM activity.

### The *scr* and *shr* mutants have an elevated level of reactive oxygen species

Because the *scr* mutant is hypersensitive to sugar (Cui et al., 2012), next we examined the effects of sugar-signaling mutants on root growth of the *scr* mutant. Double mutants were generated between *scr-1 or scr-4* and the sugar-insensitive mutants *gin2-1/hxk1-1*, *rgs1-2*, or *abi4-104* (Rolland et al., 2006). Only the *rgs1-2* mutation made the *scr* root grow longer, but to a lesser extent than the *aba2-1* mutant (Fig S2). The *scr-1 hxk1-1* and *scr-4 abi4-104* double mutants even had shorter roots than the *scr* single mutants (Fig S2). These results suggest that other processes are also altered in the *scr* mutant.

Since ROS accumulates under abiotic stress, we reasoned that redox homeostasis might be compromised in the *scr* mutants. To test this hypothesis, we compared the level of hydrogen peroxide, a major type of ROS, in the *scr* mutant and wild type roots using the DAB staining method. As shown in Fig 2, the *scr* mutant root clearly had a higher level of hydrogen peroxide in the elongation and maturation zones. The *shr* mutant root also had an elevated level of hydrogen peroxide in the elongation and maturation zones, but to a lesser extent (Fig 2). These results agree with the conclusion that SCR plays a major role in redox homeostasis.

**Fig 2.**
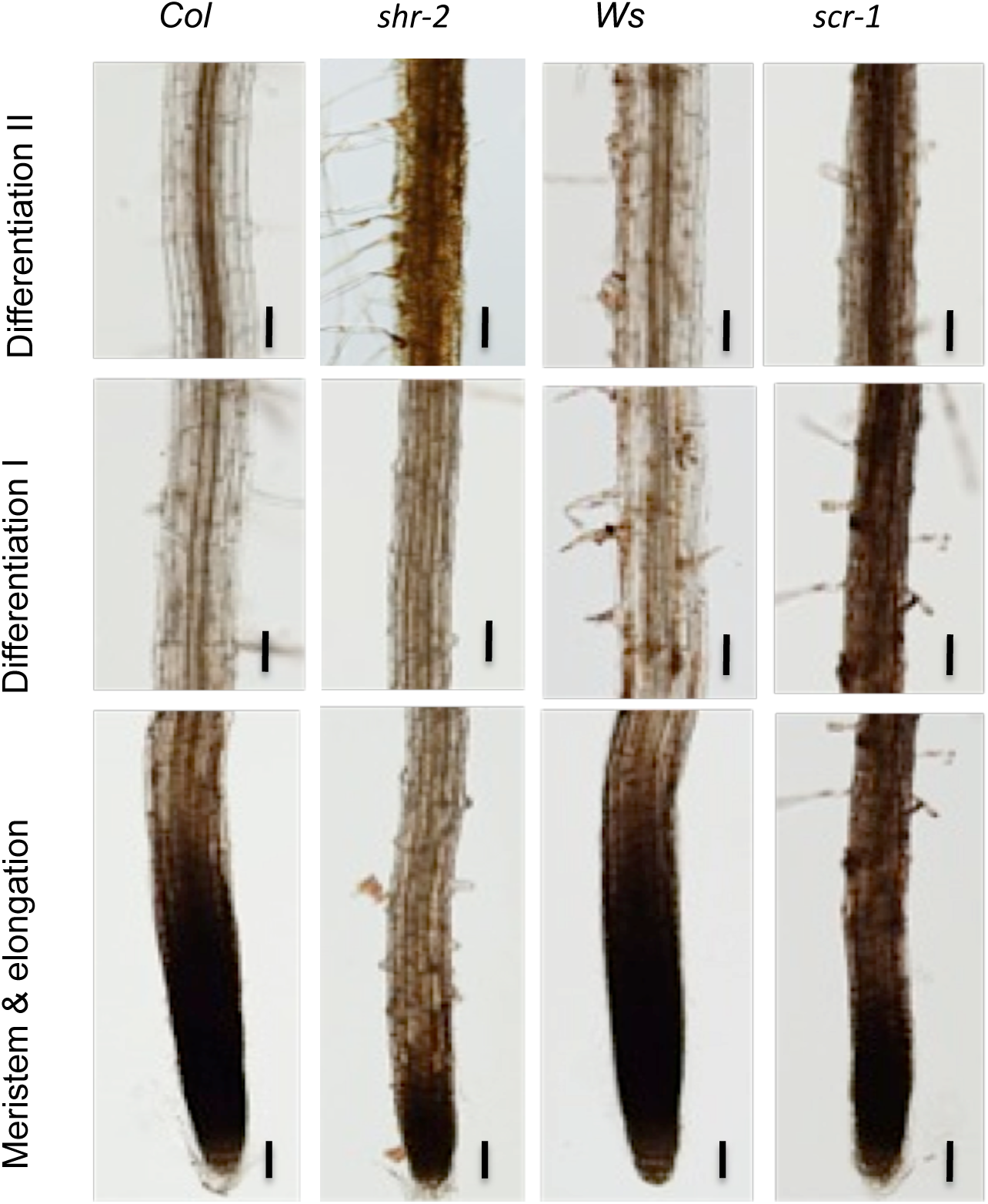
The *shr* and *scr* mutant roots have an elevated level of hydrogen peroxide. DAB staining was used to detect hydrogen peroxide in 8-day old seedlings. Bars = 50µm.

### The *aba2* mutant improves root growth in the *shr* and *scr* mutants by promoting cell elongation

Although ROS inhibit cell division and elongation, our finding that the change in ROS level in the *shr* and *scr* mutants occurs in the elongation zone, not in the meristem, raises the interesting possibility that their short-root phenotypes are also due to defect in cell elongation. We therefore measured the length of fully differentiated epidermal cells in the maturation zone. As expected, the result showed that the cells in the *shr-2* and *scr-1* mutants were dramatically shorter than those in the wild type (Fig 3A). Notably, the cells in the *scr-1 aba2-1* double mutant were significantly longer than those in the *scr-1* single mutant (Fig 3B). The cell length was also increased in the *shr-2 aba2-1* double mutant, albeit to a lesser extent (Fig 3C). Based on these results, we conclude that the *aba2* mutant improves root growth in the *shr* and *scr* mutant backgrounds not only by promoting mitotic activity but also by enhancing cell elongation. Because of the more prominent effect of *aba2* on the *scr* mutant, we have focused on SCR in the rest of this study.

**Fig 3.**
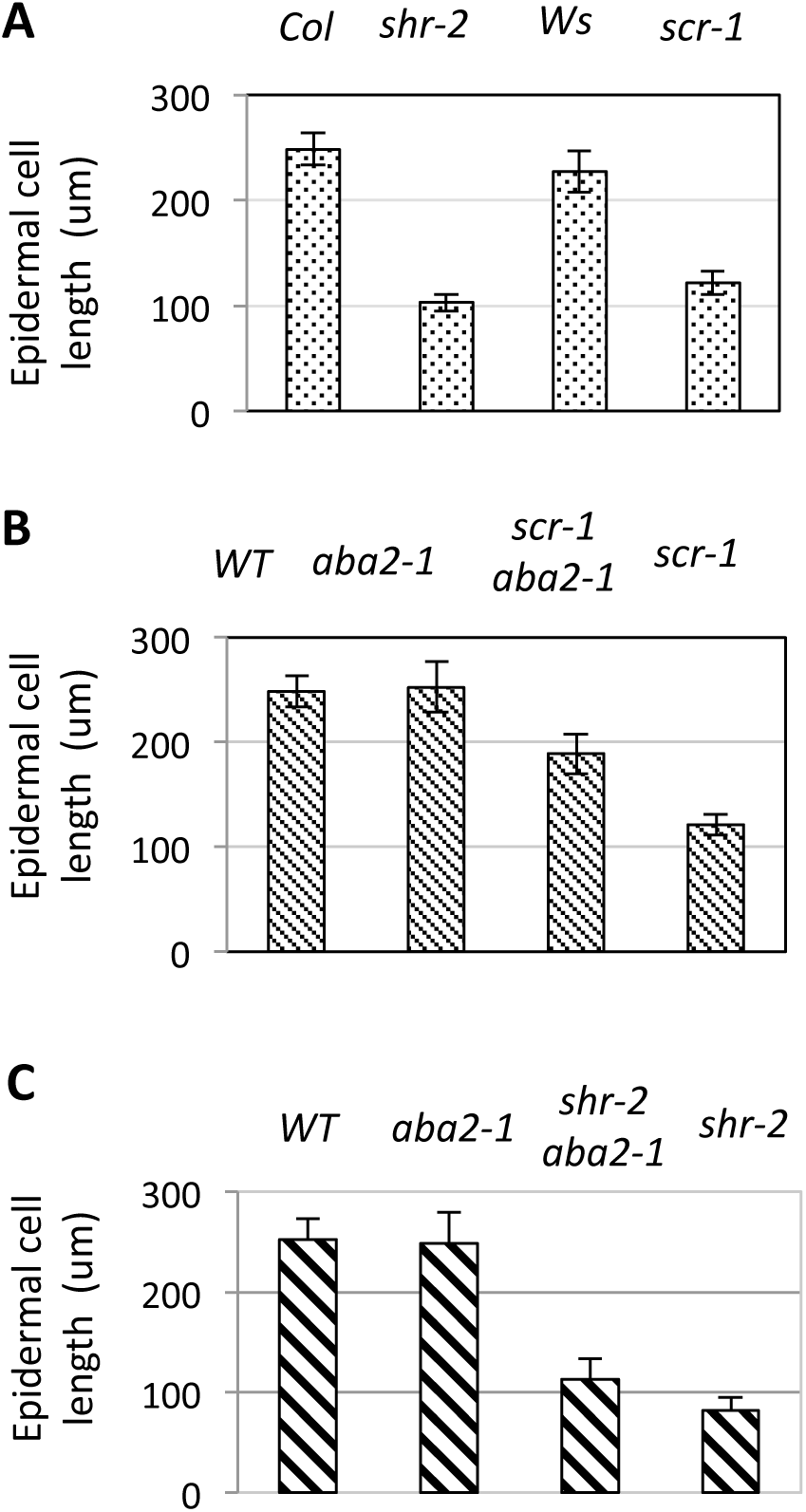
Blocking ABA signaling partially rescues the short root cell phenotype in the *shr* and *scr* mutants. A. Epidermal cell length in *shr-2* and *scr-1* mutants, and corresponding wild type (WT). Data are represented as mean ± SEM. N>20. B and C. Epidermal cell length increases in the *scr-1 aba2-1* and *shr-2 aba2-1* double mutants. Data are represented as mean ± SEM. N>20.

### The *aba2* mutation confers tolerance to oxidative stress in the *scr* mutant, but does not eliminate ROS

The *aba2-1* mutant could promote root growth by eliminating ROS or blocking oxidative stress response, or both. To distinguish among these possibilities, we first treated the *scr-1 aba2-1* double mutant, the *scr-1* and *aba2-1* single mutants, and wild type seedlings with 1mM hydrogen peroxide, which inhibits root growth but does not cause cell death according to our previous studies (Cui et al., 2014b). As shown in Fig 4A and C, relative to the *scr-1* single mutant, the *scr-1 aba2-1* double mutant had significantly longer roots, suggesting that the *aba2-1* mutation confers tolerance to oxidative stress. Next, we examined root growth in medium containing 4% glucose and found that the *aba2-1* mutation also confers the *scr* mutant tolerance to high sugar (Fig 4B and D).

**Fig 4.**
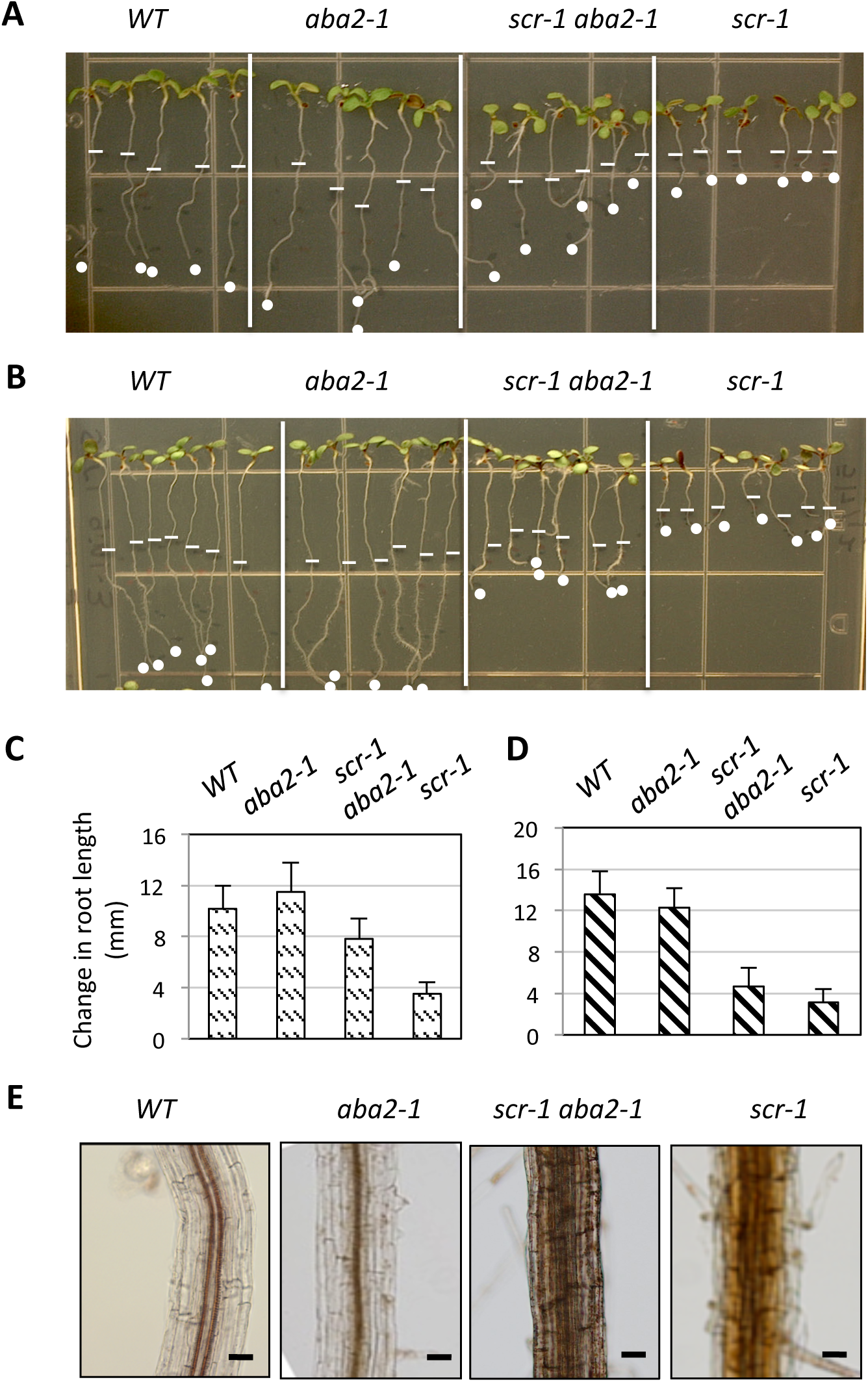
The *aba2-1* mutant confers tolerance to hydrogen peroxide and high sugar, but does not reduce hydrogen peroxide. A-B. Representative image showing root growth after seedlings (5 days old) were transferred to MS medium containing 1mM hydrogen peroxide or 4% glucose for 3 days, marked by the ticks (time of transfer) and dots (end of the treatment). C-D. Quantitative analysis of changes in root length after treatments with hydrogen peroxide or glucose, respectively. Data are represented as mean ± SEM. N > 20. E. DAB staining showing the amount of hydrogen peroxide.

Since the *scr-1 rgs1-2* double mutant also showed improved root growth, it is possible that the *rgs1-2* mutation makes the *scr* mutant more tolerant to hydrogen peroxide and high sugar as well. Although the *scr-1 rgs1-2* double mutant was slightly more tolerant to 4% sugar, it was as sensitive to 1mM hydrogen peroxide as the *scr-1* single mutant (Fig S3). This result implies that sugar stress is a minor factor affecting root growth in the *scr* mutant.

To determine whether the *aba2-1* mutant blocks ROS formation, we performed DAB staining for hydrogen peroxide in the *scr-1 aba2-1* double mutant, the *scr-1* and *aba2-1* single mutants, and wild type. No difference in hydrogen peroxide content was observed between *scr-1 aba2-1* double mutant and the *scr-1* single mutant, and between the *aba2-1* mutant and the wild type (Fig 4E). Based on the results in this section, we propose that the *aba2-1* mutant alleviates the root growth defect in the *scr* mutant mainly by mitigating the deleterious effect of oxidative stress.

### The *scr* mutant root growth defect is partially rescued by the *upb1* mutant

If the short root phenotype of *scr-1* is caused by oxidative stress, ameliorating the cellular redox status or mitigating oxidative stress response should alleviate the root growth phenotype of the *scr* mutant. To test this idea, we introduced the *upb1-1* mutation into the *scr-1* mutant, because this mutant has longer roots owing to a more reduced redox status in the root tip (Tsukagoshi et al., 2010). Although the *upb1-1* mutation had no apparent effect on root length in our growth conditions, the *scr-1 upb1-1* double mutant had significantly longer roots than the *scr-1* single mutant, a result that can be ascribed to a larger apical meristem and longer differentiated epidermal cells (Fig 5). Like the *scr-1 aba2-1* double mutant, however, the *scr-1 upb1-1* double mutant still had the stem cell and radial patterning defects characteristic of *scr*. This result lends further support to the notion that SCR promotes root growth by suppressing the oxidative stress response.

**Fig 5.**
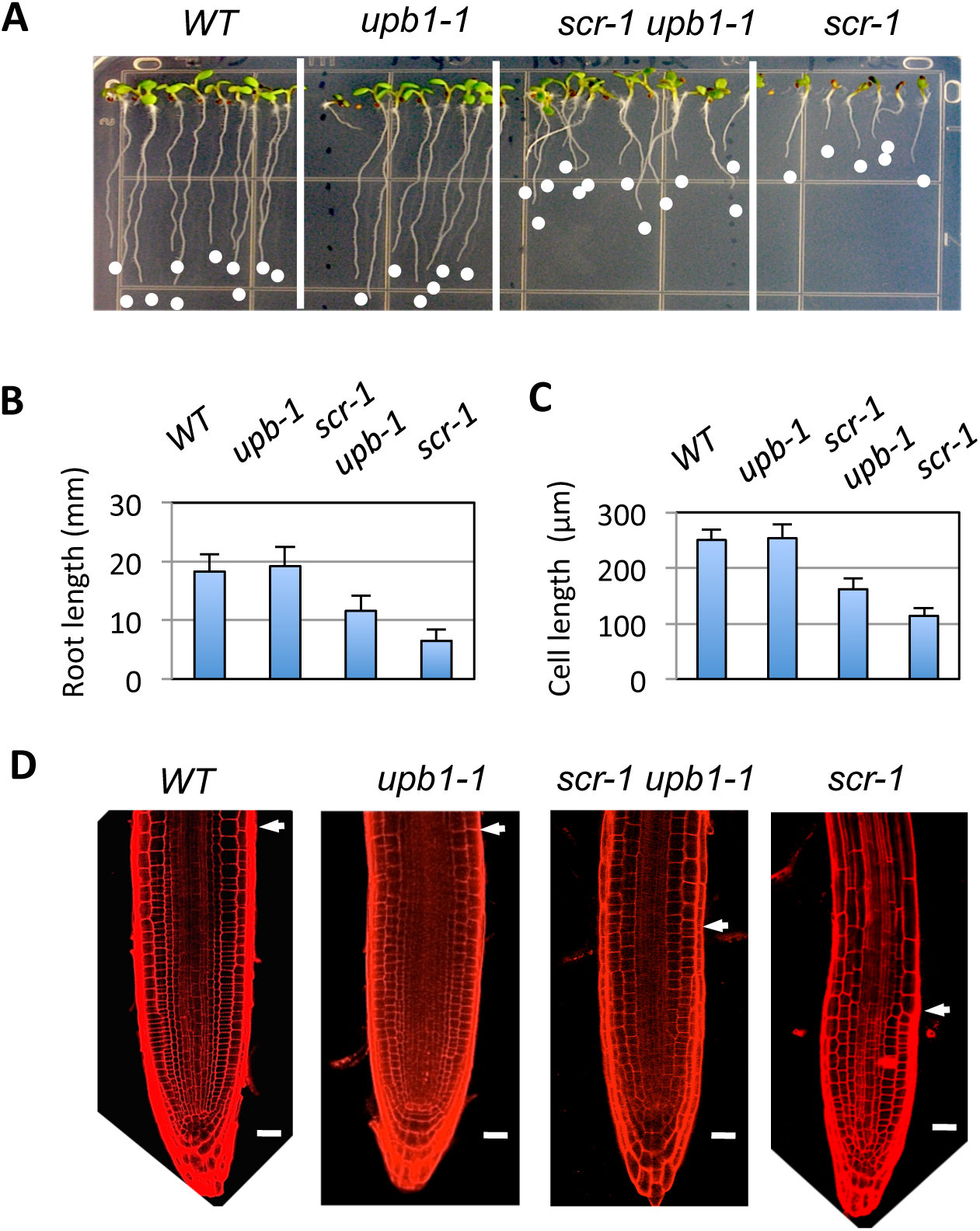
Root growth defect in *scr* mutant was partially rescued by the *upb1-1* mutant. A. Representative image of seedlings 8 days after germination. B-C. Root length and epidermal cell length in the roots of wild type, *the upb1* and *scr-1* single mutants, and the *scr-1 upb1* double mutants. Data are represented as mean ± SEM. N>20 C. Confocal microscopy image of the root rip showing the meristem size and radial patterning. Bars = 20 µm.

### Peroxidase genes are down-regulated in the *scr* mutant

Peroxidases are an important component of the cellular machinery regulating redox homeostasis. Hence, alteration in their expression could cause ROS accumulation. In the endodermis, several peroxidase (PER) genes are highly expressed, such as PER3, 9, 39, 64 and 72 (Lee et al., 2013), which are essential to Casparian strip formation in the endodermis. While converting phenolic compounds into lignin, peroxidase consumes hydrogen peroxide that is generated by NADPH oxidase (Lee et al., 2013), thus maintaining redox homeostasis. Since SCR and SHR are master regulators of endodermal differentiation (Helariutta et al., 2000; Long et al., 2015), it is likely that these peroxidase genes are down-regulated in *scr* and *shr* mutants, which would explain the elevated level of hydrogen peroxide in these mutants. To examine this possibility, we first determined the expression pattern of these peroxidase genes in the root tip of wild-type seedlings using previously reported transgenic lines that express the GFP reporter gene under the control of their promoter sequences (Lee et al., 2013). Based on the GFP fluorescence, PER3 appears to be the only peroxidase gene expressed in the elongation zone (Fig 6 and Fig S4). We therefore next compared its expression in the wild type, the *scr* mutant, and the *shr* mutant. As expected, PER3 showed a dramatic reduction in its expression in both the *shr* and *scr* mutants (Fig 6), suggesting that the elevated level of hydrogen peroxide in the *shr* and *scr* mutants is at least partially attributable to down-regulation of PER3.

**Fig 6.**
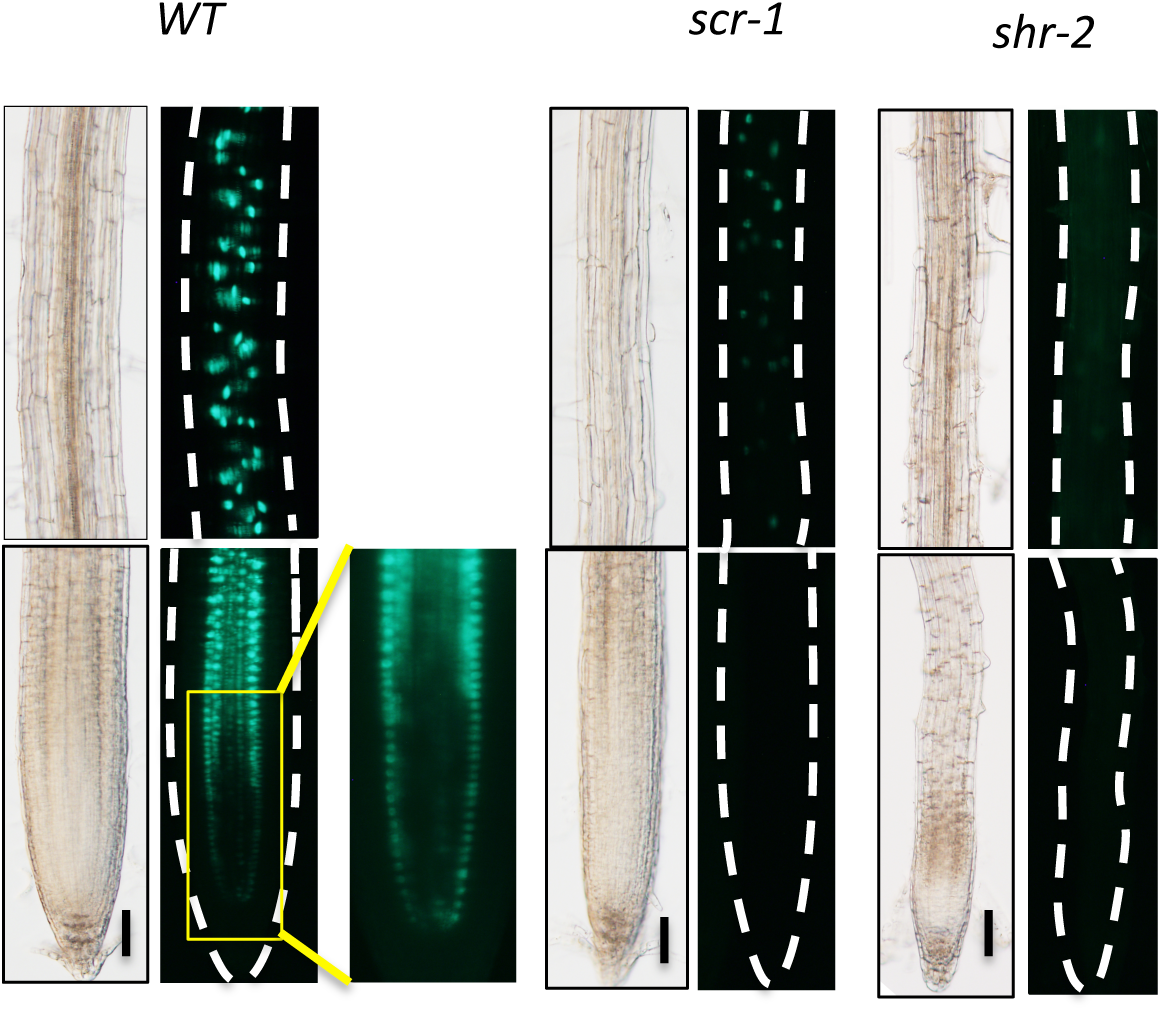
*PER3* expression level is dramatically reduced in the *scr-1* and *shr-2* mutant roots. The expression pattern of *PER3* was analyzed using the *PER3pro:nls-GFP* reporter construct. The broken lines indicate the contour of the roots. Bars = 50 µm.

### SCR directly suppresses genes involved in oxidative stress response

Our finding that the *aba2-1* mutation was able to improve root growth of the *scr* mutant without rescuing the redox defect suggests that SCR may play an active role in suppressing the oxidative stress response. In agreement with this notion, among the genes directly repressed by SCR (Cui et al., 2012; Iyer-Pascuzzi et al., 2011), we noticed that some are associated with oxidative stress. One of these genes is *WRKY15*, which is well known to be an oxidative stress responsive gene (Inze et al., 2012) and whose overexpression has been shown to cause hypersensitivity of the plant to drought (Vanderauwera et al., 2012). According to the cell-type specific RNA-seq dataset (Li et al., 2016), *WRKY15* is preferentially expressed in the endodermis (Fig S6), making it more likely a direct target of SCR. We therefore performed ChIP-PCR on the *WRKY15* promoter using transgenic plants that express a functional SCR-GFP fusion protein under its endogenous promoter, as previously described (Cui et al., 2014a; Cui et al., 2007). As shown in Fig 7A, SCR binds to a region proximal to the transcription start site of the *WRKY15* promoter (Fig 7A).

**Fig 7.**
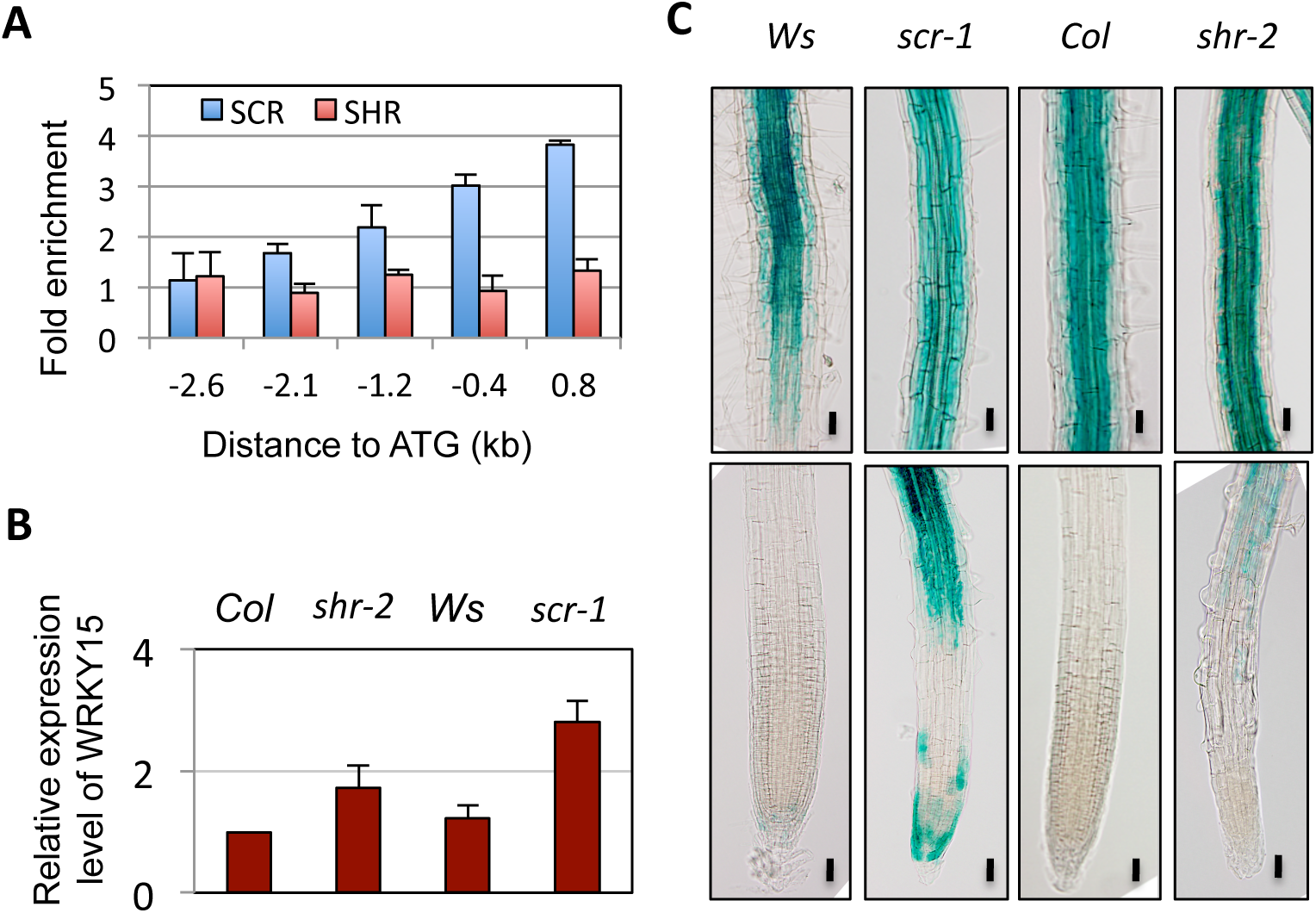
*WRKY15* is repressed directly by SCR, but not SHR. A. ChIP-PCR assay showing the binding of SCR, not SHR, to the *WRKY1*5 gene promoter. B. RT-PCR assay showing the *WRKY15* transcript level. C. *WRKY15* expression manifested using the *WRKY15pro:GUS* reporter construct. Bars = 50 μm

To determine whether *WRKY15* is suppressed by SCR, next we checked *WRKY15* transcript level in the roots of the *shr-2* and *scr*-1 mutants, as well as the wild type, by RT-PCR. Relative to the wild type, both mutants were found to have a large increase in the transcript level of *WRKY15* (Fig 7B). Notably, the change was much more pronounced in the *scr* mutant than in the *shr* mutant. To confirm this observation, next we analyzed the expression pattern of *WRKY15* using the *WRKY15*pro:GUS reporter construct, which would allow us to address the question of whether the change in *WRKY15* expression occurs in the RAM. Consistent with the RT-PCR result, *WRKY15* showed at a much higher level in the *scr* mutant (Fig 7B). Interestingly, the change in *WRKY15* expression was seen mainly in the elongation zone in both mutants (Fig 7B).

The greater prominence of SCR in regulating *WRKY15* expression suggests that SCR’s role in regulating *WRKY15* expression may be SHR-independent (Fig 7A), similar to the case with ABI4 (Cui et al., 2012). We therefore examined SHR binding to the *WRKY15* promoter by ChIP-PCR similarly as we did with SCR, but using the SHR-GFP fusion protein (Cui et al., 2011; Cui et al., 2007). No apparent binding was detected, indicating that *WRKY15* is not a direct target of SHR (Fig 7A). The results described above together suggest that SCR promote root growth by directly actively mitigating the deleterious effect of oxidative response.

## Discussion

### SCR promotes root growth through multiple mechanisms

It has been over two decades since SCR was first identified as a key regulator of root growth and development, but only recently have the underlying mechanisms begun to be elucidated. In the *scr* mutant, Moubayidin *et al.* found that some cytokinin response genes are elevated (Moubayidin et al., 2013). In further studies, Moubayidin *et al.* showed that the meristem defect in the *scr* mutant could be rescued by mutations in some genes involved in cytokinin signaling, which led them to conclude that SCR promotes root growth by suppressing the cytokinin response (Moubayidin et al., 2013). The rescue is partial, however, suggesting the existence of other mechanisms. Consistent with this, there is evidence that SCR expressed in the elongation and maturation zones is also required for root growth (Moubayidin et al., 2016; Sabatini et al., 2003). In the present study, we showed that this role of SCR can be attributed to its function as a regulator of redox homeostasis and oxidative stress response. Supporting this conclusion, we showed that the *scr* mutant had an elevated level of hydrogen peroxide in the elongation region, but not in the meristem zone, and that the root growth defect could be partly rescued when abiotic stress response was blocked using the ABA signaling mutant *aba2-1* or when cellular redox status was ameliorated using the redox regulator mutant *upb1-1*. Hence, it seems that SCR promotes root growth at least through two distinct mechanisms: one by repressing cytokinin response in the meristem, and the other by maintaining cellular redox homeostasis and mitigating the oxidative stress response in the elongation zone.

### SCR promote root growth by enhancing cell elongation and mitotic activity

It is generally accepted that SCR and SHR promote root growth by maintaining the QC. However, this view has been challenged by recent findings that the root growth defect in the *shr* and *scr* mutants can be largely rescued by mutants deficient in cytokinin biosynthesis or signaling while the QC defect still persists (Moubayidin et al., 2013; Sebastian et al., 2015). In the present study, we also found a QC-independent root growth phenomenon – although the *scr aba2* double mutant had a significantly longer root than the *scr* single mutant, it still had the QC and radial patterning defects.

Cell elongation plays a role, equally significant, if not more, than cell division in plant growth, but the mechanisms regulating root cell elongation have been unclear. In this study we found that SHR and SCR have a previously unrecognized role in promoting cell elongation, which can be attributed to their function in regulating redox homeostasis and oxidative stress response. Using differentiated epidermal cells as a proxy, we first showed that cells in the *shr* and *scr* mutant roots were dramatically shorter than those in the wild type. By DAB staining we found that the level of hydrogen peroxide was significantly elevated in the *shr* and *scr* mutant roots, and this change was seen in the elongation zone but not in the meristem zone. Consistent with this observation, PER3, an enzyme that consumes hydrogen peroxide for the formation of the Casparian strip, almost completely lost its expression in the elongation zone, whereas *WRKY15*, an oxidative stress response gene, was up-regulated. Last but not least importantly, the short cell defect in the *scr* root was largely rescued by the *aba2-1* and *upb1-1* mutants, which block the abiotic stress response or ameliorate the redox status respectively.

Interestingly, the *scr-1 aba2-1* and *scr-1 upb1-1* double mutant roots also had larger and mitotically more active apical meristems, suggesting that SCR may also promote meristem activity through its role in redox homeostasis and oxidative stress response. Since there was no apparent change in redox status in the meristem zone in the *scr* mutant, we think that this effect is an indirect consequence of a prolonged phase of cell elongation resulting from an improved growth microenvironment. Many hormones and environmental factors are known to affect the meristem size by promoting cell differentiation (Del Pozo, 2016; Moubayidin et al., 2013). By extending the duration of cell elongation, SCR could slow down the cell differentiation program, thereby helping to maintain a larger meristem.

### The role for SCR in redox homeostasis and oxidative stress response is independent of SHR

Although SCR acts downstream of SHR in stem cell renewal and radial patterning, our results indicate that SCR plays a more prominent role in redox homeostasis and oxidative stress response. First, relative to the *shr* mutant, the *scr* mutant appears to have more ROS. Second, although the root growth defect of the *shr* mutant was partially rescued by the ABA signaling mutant *aba2-1*, the effect is less pronounced. Notably, the root growth defect was even not rescued by the *upb1-1* mutation. Furthermore, ChIP-PCR assay showed that SCR but not SHR binds to the promoter of *WRKY15*. This SHR-independent role of SCR is not unexpected, as previously we have shown that ABI4 is also directly regulated by SCR, but not SHR (Cui and Benfey, 2009a). On the other hand, SHR also has an SCR-independent role in cytokinin homeostasis and radial patterning in the stele (Cui et al., 2011; Levesque et al., 2006). Consistent with this, blocking cytokinin signaling in the *shr* mutant has been shown to be able to restore root growth to the wild-type level, but this only partially rescues the root growth defect in the *scr* mutant, even though the root growth phenotype is more severe in the *shr* mutant (Sebastian et al., 2015). The different functions of SHR and SCR can be ascribed partly to their different expression domains and partly to SCR proteins that are not bound by SHR resulting from a feedback mechanism for SCR expression.

### SCR is a key player coordinating cell elongation and endodermal differentiation

ROS are inhibitory to cell elongation and division and therefore must be removed to ensure normal growth and development. The oxidative stress response must also be suppressed, as it is mounted at the expense of developmental processes. ROS, however, are essential to endodermal differentiation owing to their positive role in Casparian strip formation. SCR resolves this conflict by the repressing oxidative stress response while activating peroxidase expression. This study has thus revealed an important mechanism coordinating multiple processes in root growth and development.

## Materials and Methods

### Plant materials and treatments

Double mutants were generated by crossing and confirmed by PCR-based genotyping. If the two mutants had genetic backgrounds (Col, Ws or Ler), then mutants as well as wild type in the F2 segregating population were distinguished based on their growth type and seeds in the genetic background were used in the same comparisons. Seeds were surface-sterilized for 5-10 minutes using 20% bleach plus 0.1% Tween 20, then washed 4 times with sterile H_2_O. After storage at 4°C for 1 week for vernalization, the seeds were sowed on growth medium containing 1X MS, 0.8% agar (RPI, Cat No. A20400) and 1% sucrose and allowed to germinate in a vertical position in a Percival→ growth chamber at 22°C with 16 h daily illumination. For sugar tolerance assay, glucose was added to the standard MS medium to a final concentration of 4%, whereas for hydrogen peroxide treatment 2ml of 1mM hydrogen peroxide solution was added to the surface of the MS growth medium.

### Histochemical staining

GUS staining was performed essentially as described in the Arabidopsis book (Weigel and Glazebrook, 2002). Seedlings were incubated in GUS staining solution overnight at 37°C. For hydrogen peroxide detection, the DAB staining protocol by (Daudi and O’Brien, 2012) was followed. To visualize starch granules, the roots were incubated in Lugol solution (Sigma, L-62650) for half an hour at room temperature (RT), followed by washing with distilled water.

### Microscopy

For light microscopy, the roots were first cleared in a drop of chloral hydrate solution (7.5 g chloral hydrate dissolved in 3 ml 50% glycerol) on a glass slide for 1-5 minutes. Images were captured using an Olympus BX61 compound microscope. For confocal microscopy, the seedlings were stained in Propidium iodide (Sigma, P-4170) solution (5 µg/ml in ddH_2_O) for 1 minute at RT and imaged with a Zeiss Meta510 confocal microscope.

We define the root meristem as the region between the QC and the cells that are two times longer than the meristem cells. To measure the meristem size, we first cleared the roots in chloral hydrate solution, imaged them with an Olympus BX61 compound microscope, and then counted the number of cortex cells in a single file. The epidermal cell length was measured similarly, but with the use of the Image J software (https://imagej.nih.gov/ij/).

### Molecular biology assays

Chromatin immunoprecipitation (ChIP)) was performed as we previously described (Cui et al., 2007), using the GFP antibody (ab290, ABcam, MA) and transgenic plants expressing the SCR-GFP and SHR-GFP fusion proteins under their endogenous promoters in the *scr-1* or *shr-2 mutant* backgrounds. As a negative control, a mock ChIP was also conducted whereby BSA serum was used instead of the antibody. Quantitative PCR was then performed on the *WRKY15* promoter using an ABI 7000 real-time thermocycler, with the 18S rDNA as a loading control. The primers used for these experiments are shown in Table S1.

For RT-PCR, 1 µg total RNA isolated with the RNeasy Plant Mini Kit (Qiagen) was first converted into cDNA using the Superscript III First Strand Synthesis System (Invitrogen). Quantitative PCR analysis was then conducted using the ABI 7000 real-time thermocycler, similar to the ChIP-PCR using gene-specific primers (Table S1). The 18S rRNA was used as a control for the amount of RNA and reverse transcription.

The CTAB method was used for preparation of genomic DNA used in genotyping and the primers used for this purpose are listed in Table S1.

## Acknowledgements

The authors are thankful to Dr. Niko Geldner (University of Lausanne, Switzerland) for the PERpro:GFP transgenic lines, Dr. Frank Van Breusegem (Ghent University, Belgium) for the *WRKY15*pro:GUS transgenic line, Dr. Philip Benfey (Duke University) for the *upb1-1* mutant, and Dr. Jen Sheen (Harvard University, USA) for the *hxk1-1* mutant. The *rgs1-2* (SALK_074376), *abi4-104* and *aba2-1* mutants were obtained from the Arabidopsis stock center. The authors are also thankful to Jen Kennedy (Florida State University) for editing the manuscript. This research is supported by the Florida State University, the Northwest Agriculture and Forest University, and the National Science Foundation of China (grant No. 31871493). Imaging was conducted at the Biological Science Imaging Core Facility of the Florida State University.

